# Patient satisfaction with nursing care in Ethiopia: A systematic review and meta-analysis

**DOI:** 10.1101/544783

**Authors:** Henok Mulugeta, Fasil Wagnew, Getenet Dessie, Henok Biresaw, Tesfa Dejenie Habtewold

**Affiliations:** Lecturer of Nursing, Department of Nursing, College of Health Science, Debre Markos University Address: P.O. Box 269, Debre Markos, Ethiopia; Lecturer of Nursing, Department of Nursing, School of Health Science, College of Medicine and Health Science, Bahir Dar University Address: P.O. Box 79, Bahir Dar, Ethiopia; Lecturer of Nursing, Department of Nursing, College of Health Science, University of Gondar Address: P.O. Box 196, Gondar, Ethiopia; Department of Epidemiology and psychiatry, University of Groningen, University Medical Center Groningen, Groningen, Netherlands. Department of Nursing, Debre Birhan University, Debre Birhan, Ethiopia Address: P.O. Box 445, Debre Berhan, Ethiopia

**Keywords:** Nursing care, patient satisfaction, systematic review, meta-analysis, Ethiopia

## Abstract

**Background:** Patient satisfaction with nursing care has been considered as the most important predictor of the overall patient satisfaction with hospital service and quality of health care service at large. However, the national level of patient satisfaction with nursing care remains unknown. Hence, the objective of this systematic review and meta-analysis was to estimate patient satisfaction with nursing care in Ethiopia.

**Methods:** Studies were accessed through an electronic web-based search strategy from PubMed, Cochrane Library, Google Scholar, Embase, PsycINFO and CINAHL by using combination search terms. Qualities of each included article assessed by using a modified version of the Newcastle-Ottawa Scale for cross-sectional studies. All statistical analyses were done using STATA version 14 software. The Preferred Reporting Items for Systematic Reviews and Meta-Analyses (PRISMA) guideline was followed for reporting results.

**Results:** Of 1,166 records screened, 15 studies with 6091 participants were included. The estimated pooled level of patient satisfaction with nursing care in Ethiopia was 55.15% (95% CI (47.35, 62.95%)). Based on the subgroup analysis, the estimated level of patient satisfaction was 61.84% (95% CI: 44.49, 79.2) in Addis Ababa, 54.24 %(95%CI: 46.84, 61.65) in Amhara region, 44.06% (95%CI: 38.09, 50.03) in SNNP, and 53.02 %(95% CI: 50.03, 56.00) in other regions. Patients who have one nurse in charge [(OR 1.08(0.45, 2.62)], with no history of previous hospitalization [(OR 1.37(0.82, 2.31)], living in the urban area [(OR 1.07(0.70, 1.65)], / and those who have no comorbid disease [(OR 1.08(0.48, 2.62)] were more likely to be satisfied with nursing care than their counterparts even though it was not statistically significant.

**Conclusion:** This meta-analysis revealed that about one in two patients were not satisfied with the nursing care provided in Ethiopia. Therefore, Ministry of Health should give more emphasis to the quality of nursing care in order to increase patient satisfaction which is important to improve the overall quality of healthcare service.

## Background

Quality healthcare delivery and creation of patient satisfaction are the primary goals of hospitals[1]. One of the ways of evaluating the performance of health care is assessing patient satisfaction with the nursing care since it was considered as a fundamental indicator of quality health care provided in hospitals[2, 3].

Patient satisfaction simply described as the value and reaction of patients towards the care they received[4]. Moreover, according to the American Nursing Association patient satisfaction defined as patients’ value and attitude of care they received from the nursing staffs during their hospitalization[5].

Today, patient satisfaction is the major concern of healthcare institutions. Satisfied patients are more likely to have a good relationship with the nurses which result in improved quality of care[6, 7]. Literature also suggests that patient satisfaction is directly linked to better patient outcomes. Furthermore, achieving patient satisfaction with nursing care result in better patient compliance with health care regimens and better health outcomes[8].

Nurses are a pivotal part of the health care system and they spent more time with patients. Moreover, nurses provide about 80% of primary health care service in the hospital. Hence, patient satisfaction with the nursing care can determine the overall satisfaction of the hospital service provided [8–10].

Determining the factors that influence patient satisfaction is important for nurses to improve the quality of nursing care. Patient satisfaction with nursing care can be affected by numerous factors since it is a complex and multidimensional concept[8, 11]. For example, some literature suggested that availability of an assigned nurse, behaviors of nurses, the surrounding physical environment, and history of the previous hospitalization are the major determinant factors for the overall patient satisfaction[4, 8, 12–14]. While others showed that sociodemographic factors like age, residence and educational level are the most determining factors for patient satisfaction[15–18].

In recent years, many studies have been conducted to determine the level of patient satisfaction with nursing care. For instance, a study was done in Iraq[19], Brazil[20] and Egypt[21] showed that patients were highly satisfied with the nursing care. Additionally, the overall level of patient satisfaction with nursing care is 69% in Iran[22], 67% in Kenya[23], and 33% in Ghana[24]. On the contrary, the results of the study done in India revealed that most of the hospitalized patients have poor perception regarding nursing care[25].

The Ethiopian Federal Ministry of Health is striving to provide quality health care service in the country by developing different quality management guidelines and health sector development plans in order to increase patients’ satisfaction with the healthcare service[26–28].

Though few patient satisfaction surveys with nursing care have been conducted previously in different areas of Ethiopia, the overall level of patient satisfaction with nursing care in the country level remains unknown. Moreover, they are not consistent and inconclusive to determine the level of patient satisfaction at the country level. In addition, determining the level of patient satisfaction with nursing care appears crucial to monitor and improve the quality of nursing care. Therefore, the objective of this systematic review and meta-analysis is to estimate the pooled level of patient satisfaction with nursing care and to identify the contributing factors in Ethiopia.

## Methods

### Design and search strategy

The procedure for this systematic review and meta-analysis was designed in accordance with the Preferred Reporting Items for Systematic Reviews and Meta-Analyses (PRISMA) guidelines[29]. We searched PubMed, Cochrane Library, Google Scholar, CINAHL, Embase, and PsycINFO database for studies reporting the level of patient satisfaction with nursing care from study conception to May, / 2018. EndNote (versiaon X8) reference management software was used to download, organize, review and cite the related articles. Comprehensive search was performed using the following search terms: “Patient satisfaction”, “satisfaction”, “determinants of patient satisfaction”, “nursing care”, and “Ethiopia”. “AND” and “OR”. Boolean operators were used to combine search terms. Furthermore, we manually searched cross references in order to identify additional relevant articles.

### Inclusion and Exclusion criteria

We included studies reporting patient satisfaction with nursing care among admitted patients and its determinants in irrespective of their sex and other demographic characteristics. Studies were also included if they assessed the determinants of patient satisfaction with nursing care. Both published and gray literature reported in the English language regardless of date of study/publication were also included. Nevertheless, articles without full text and with poor quality were excluded. Two authors (H.M. and G.D.) independently evaluated the eligibility of all retrieved studies, and any disagreement and inconsistencies were resolved by discussion and consensus.

### Data extraction and quality assessment

After the screening was completed, the relevant data from each included article were extracted using a pre-piloted data extraction format prepared in a Microsoft Excel spreadsheet. Data on author/s name, year of publication, study area/Region, health institution, study design, sample size, prevalence and determinant factors were extracted from each included article by three independent authors (H.M. FW, and G.D). Disagreement and inconsistencies were resolved by discussion among the authors.

The Joanna Briggs Institute Prevalence Critical Appraisal Tool for use in systematic review for prevalence study was used for critical appraisal of studies[30]. Moreover, methodological and other quality of each article was assessed based on a modified version of the Newcastle-Ottawa Scale for cross-sectional study adapted from Modesti et al[31]. Two authors (HM, HB) independently assessed the quality of each article. Whenever it is necessary a third reviewer (TDH) were consulted. Any disagreement was resolved through discussion and consensus.

### Statistical Analysis

The extracted data were transferred to STATA version 14 for meta-analysis. Meta-analysis of the level of patient satisfaction with nursing care was carried out using a random effects model, generating a pooled effect size with 95% confidence interval (CI). The effect of selected determinant variables was independently analyzed and was presented using a forest plot. We also reported measures of association using the ORs with a 95% CI. All data manipulation and statistical analyses were performed using Stata version 14.0 software.

Heterogeneity across studies was evaluated using I^2^ test statistics and Cochrane Q statistics. I^2^ statistics is used to quantify the percentage of total variation in study estimate due to heterogeneity. I^2^ ranges between 0 and 100%. I^2^ >=75% indicate very high heterogeneity across the studies. A p-value of less than 0.05 was used to declare significant heterogeneity[32, 33]. The random effects model using Der Simonian and Laird method is the most common method in a meta-analysis to adjust for the observed variability[34, 35]. Furthermore, the source of heterogeneity was also assessed by subgroup analysis based on region and meta-regression.

A Funnel plot was used for visual assessment of publication bias. Asymmetry of the funnel plot is an indicator of potential publication bias[36]. We also employed Egger’s and the Begg ‘s test to determine if there was significant publication bias. A p-value of less than 0.05 was considered significant[37]. Finally, we performed a sensitivity analysis to describe whether the pooled effect size was influenced by individual studies.

## Results

### Search result and study characteristics

The electronic online search yielded 1166 records, of which 42 duplicate records identified and removed. Title and abstract screening result in the exclusion of 1042 non-relevant articles. From the remaining 82 articles, 28 articles were excluded since they are on general hospital service. Then, 54 articles underwent for full-text screening. However, 39 articles were excluded based on our predetermined eligibility criteria. Finally, a total of 15 articles included in the meta-analysis (Figure 1).

A total of 15 studies with 6,091 participants were included in this meta-analysis. Among 15 studies five[14, 28, 38–40] were conducted in Addis Ababa, five [26, 41–44] were conducted in Amhara region, two [27, 45] were in SNNP region, and three studies[12, 46, 47] were in other regions(Oromia, Harari and Tigray). All studies were a cross-sectional study conducted among admitted adult patients in different hospitals of Ethiopia (Table 1).

Figure 1: PIRSMA Flowchart diagram of the study selection

**Table 1:** Characteristics of studies included in the meta-analysis of patient satisfaction with nursing care

| S.No | Author/s[reference] | Publication Year | Study area, Region | Health Facility Name | Study Design | Sample Size | Proportion % (95%CI) |
| --- | --- | --- | --- | --- | --- | --- | --- |
| 1 | Mulugeta M. et al [28] | 2014 | Addis Ababa | Black Lion Hospital | Cross-sectional | 374 | 90.1(87.1,93.1) |
| 2 | Getachew G. et al [38] | 2016 | Addis Ababa | Menelik Hospital | Cross-sectional | 372 | 46.7(41.6,51.8) |
| 3 | Solomon Bekele [39] | 2009 | Addis Ababa | Addis Ababa Public Hospitals | Cross-sectional | 435 | 56.3(51.6,61.0) |
| 4 | Bekele Chaka [40] | 2005 | Addis Ababa | Addis Ababa Public Hospitals | Cross-sectional | 631 | 67.0(63.3,70.7) |
| 5 | Melsew Getinet [14] | 2017 | Addis Ababa | Addis Ababa Public Hospitals | Cross-sectional | 422 | 48.8(44.0,53.6) |
| 6 | Melesse Belayneh [26] | 2016 | Bahir Dar, Amhara | Felege Hiwot Referral Hospital | Cross-sectional | 236 | 44.9(38.6,51.2) |
| 7 | Negash AK. et al [43] | 2014 | Bahir Dar, Amhara | Felege Hiwot Referral Hospital | Cross-sectional | 373 | 67.1(62.3,71.9) |
| 8 | Sharew NT. et al [44] | 2018 | Debre Birhan, Amhara | Debre Birhan Referral Hospital | Cross-sectional | 384 | 49.2(44.2,54.2) |
| 9 | Alemu S. et al [41] | 2014 | Debre Markos, Amhara | Debre Markos Referral Hospital | Cross-sectional | 392 | 56.9(52.0,61.9) |
| 10 | Haile Eyasu K. et al [42] | 2016 | Dessie, Amhara | Dessie Referral Hospital | Cross-sectional | 374 | 52.5(47.4,57.6) |
| 11 | Mensa M. et al [27] | 2017 | Arba Minch, SNNP | Arba Minch General Hospital | Cross-sectional | 236 | 40.9(35.5,46.3) |
| 12 | Legesse MT. et al [45] | 2016 | Hawassa, SNNP | Hawassa University Specialized Hospital | Cross-sectional | 406 | 47.0(42.2,51.9) |
| 13 | Jiru TG. et al [46] | 2017 | Nagele Borena, Oromia | Nagele Borena And Adola General Hospital | Cross-sectional | 413 | 55.9(51.1,60.7) |
| 14 | Ahmed T. et al [12] | 2014 | Harar, Harari | Public Hospitals in Eastern Ethiopia | Cross-sectional | 582 | 52.7(48.6,56.8) |
| 15 | MollaTeferi [47] | 2017 | Mekelle, Tigray | Ayder Specialized Hospital | Cross-sectional | 374 | 50.3(45.2,55.4) |

### Patient satisfaction with nursing care

The pooled effect size of patient satisfaction with nursing care using the fixed effect model showed a significant heterogeneity across the studies. Therefore, we performed the analysis with a random effects model with 95% CI in order to adjust for the observed variability. Using random effects model, the overall estimated pooled level of patient satisfaction with nursing care as reported by the 15 studies was 55.15% (95% CI (47.35, 62.95%)) with significant heterogeneity between studies (I^2^=97.7, P=0.001). The pooled effect size of patient satisfaction with nursing care presented using forest plot (Figure 2).

**Figure 2:**
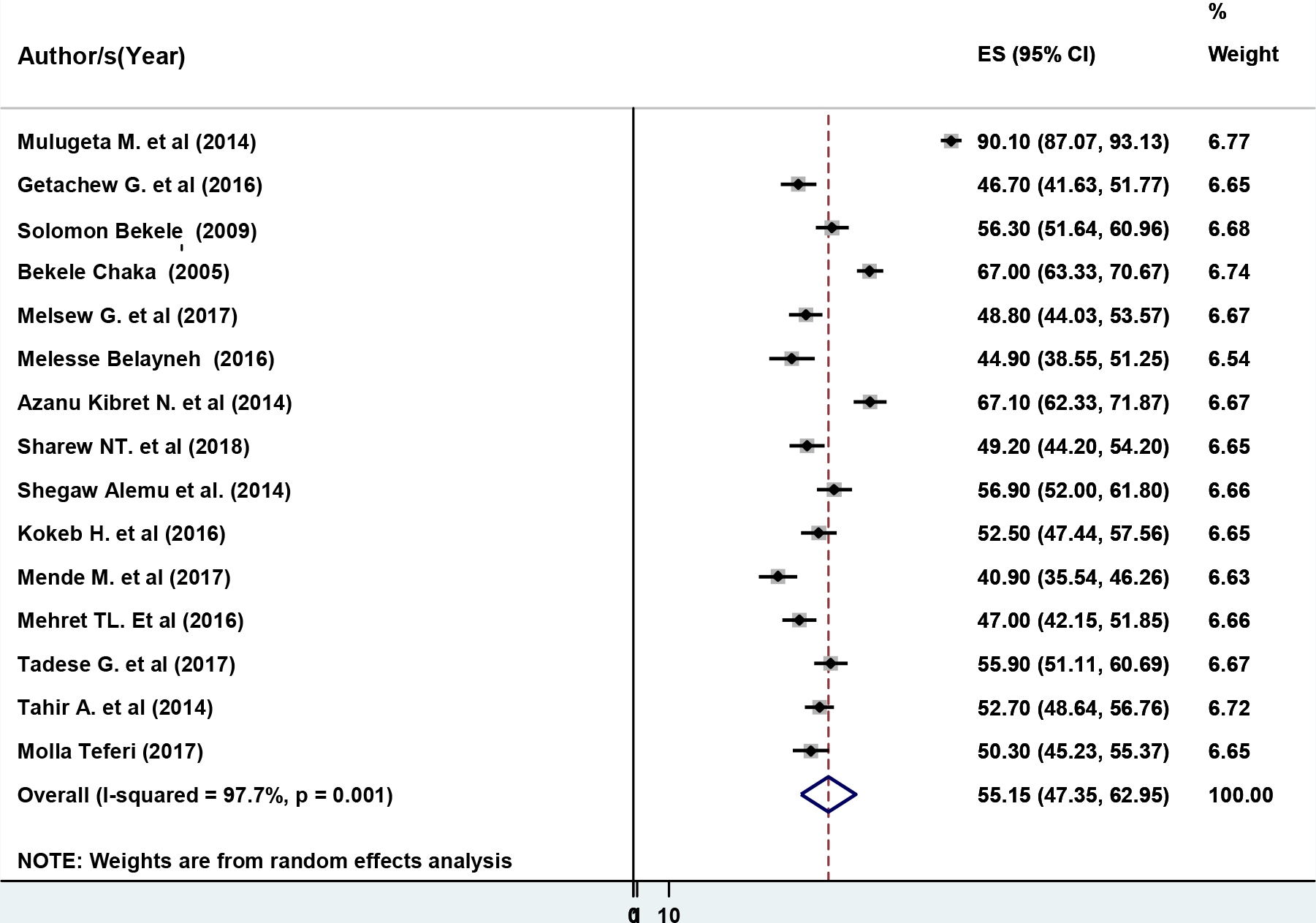
Forest plot showing the pooled level of patient satisfaction with nursing care

Subgroup analysis by region in Ethiopia was conducted to compare the level of patient satisfaction with nursing care. Based on the subgroup analysis, the highest estimated level of patient satisfaction was observed in Addis Ababa (61.84% (95% CI: 44.49, 79.2), *I^2^* = 98.9%), followed by Amhara region (54.24% (95% CI: 46.84, 61.65), *I^2^* = 90.3 %). Moreover, the lowest estimated level of patient satisfaction was observed in SNNP region (44.06% (95% CI: 38.09, 50.03), I^2^= 63.4%) (Figure 3).

**Figure 3:**
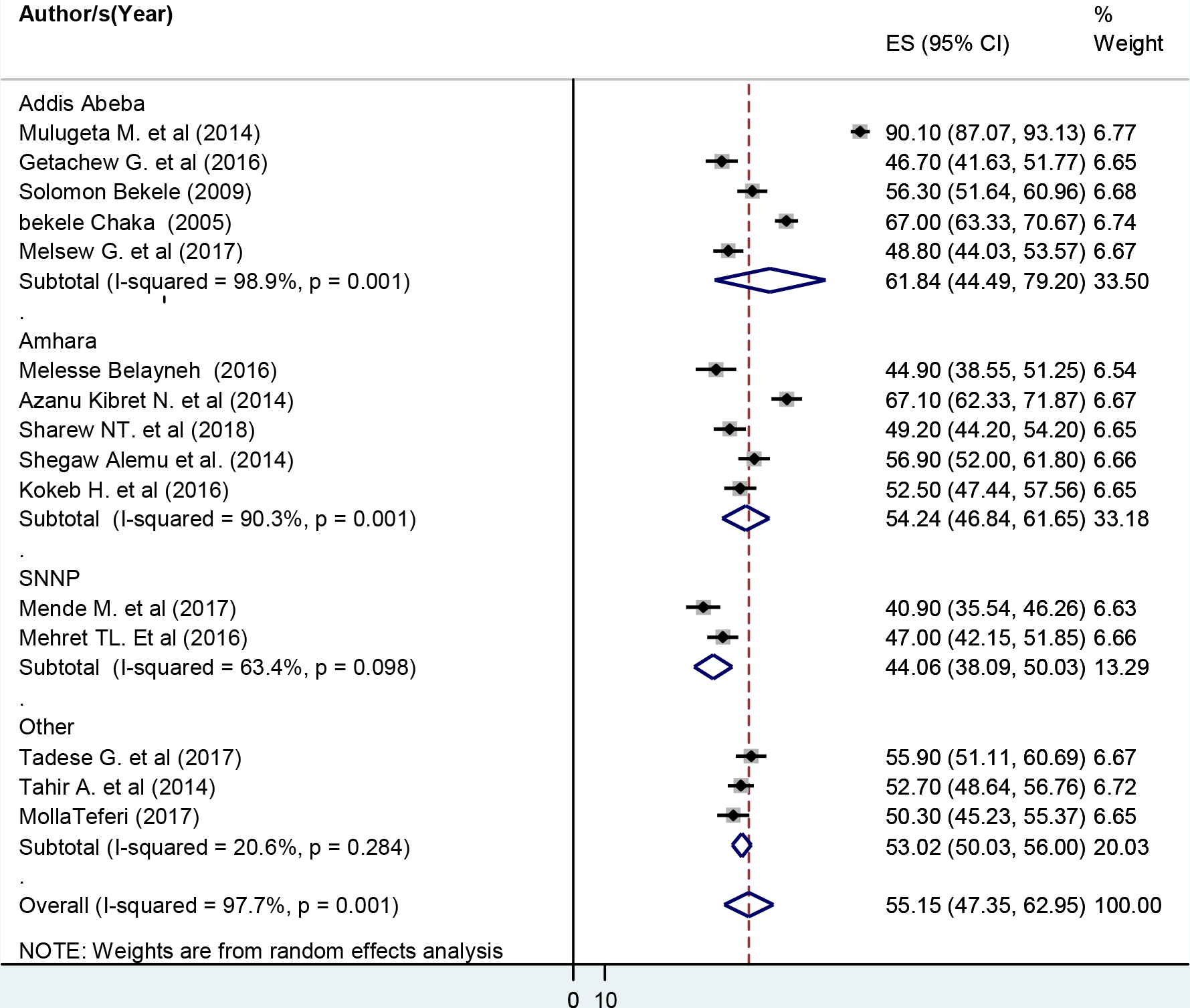
Subgroup analysis by regions on the level of patient satisfaction with nursing care

### Investigation of heterogeneity and publication bias

Given that the result of this meta-analysis revealed a statistically significant heterogeneity among studies (I^2^ statistics=97.7%), we performed subgroup analysis by region in order to minimize heterogeneity (Figure 3). Furthermore, to identify the possible source of heterogeneity, we performed meta-regression using sample size and publication year as covariates. However, the result of the meta-regression analysis showed that both covariates were not statistically significant for the presence of heterogeneity (Table 2).

**Table 2:**
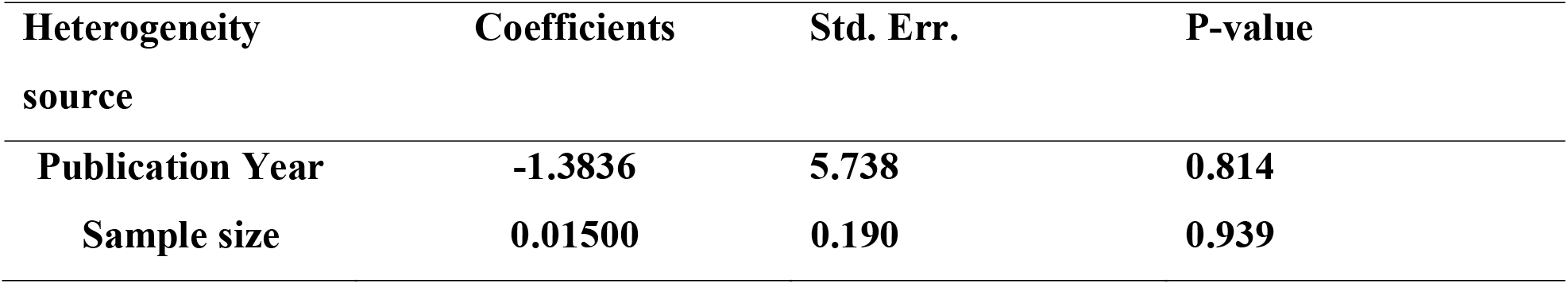
Meta-regression analysis of factors with heterogeneity of patient satisfaction with nursing care in Ethiopia, 2018.

Presence of publication bias was examined using funnel plot and Egger’s test. Visual inspection of the funnel plot suggests symmetry (Figure 4). However, the result of Egger’s test is statistically significant for the presence of publication bias (p=0.001). Moreover, the result of sensitivity analyses using random effects model suggested that no single study influenced the overall estimate (Figure 5).

**Figure 4:**
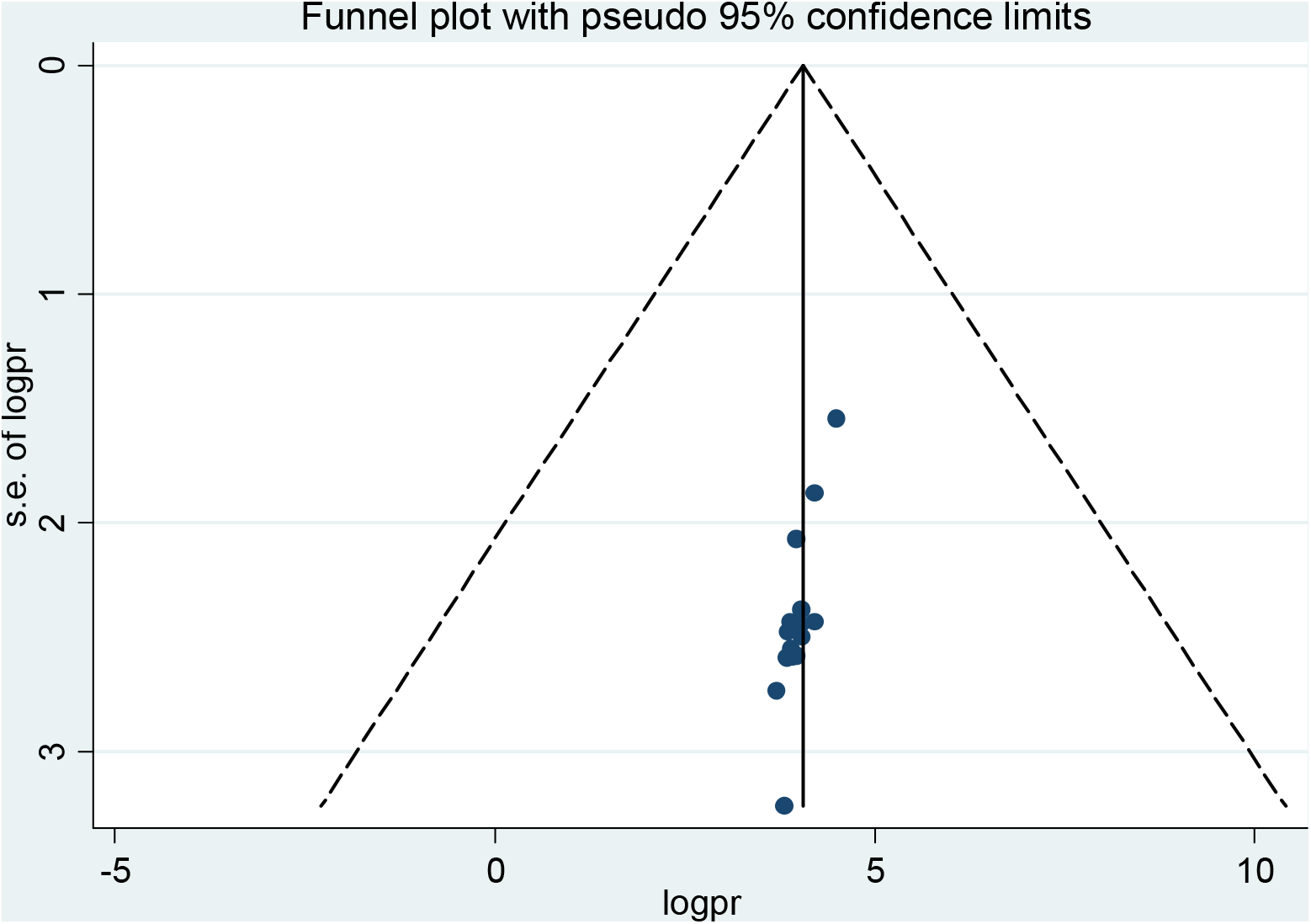
Funnel plot to test publication bias of the 15 studies

**Figure 5:**
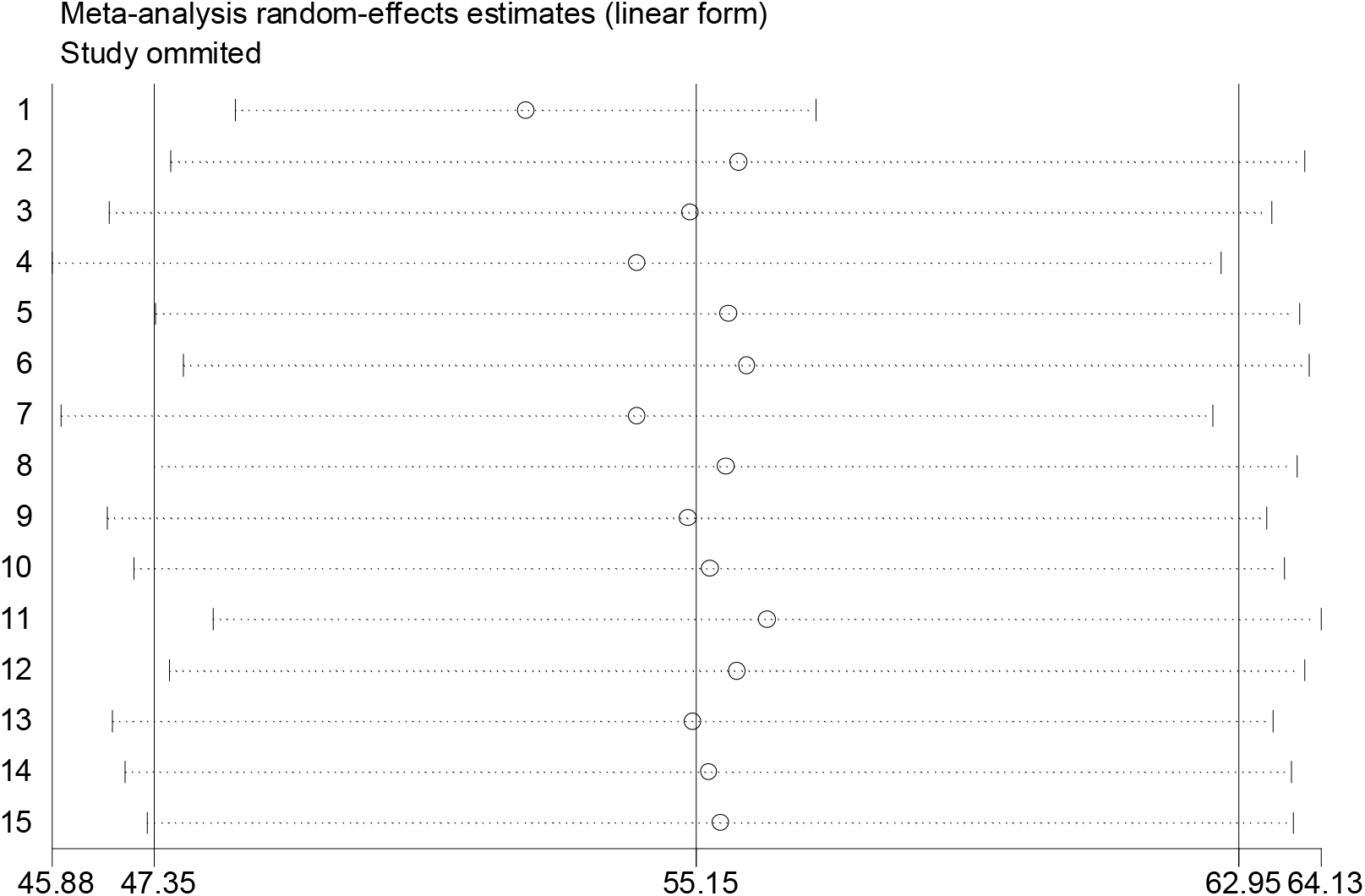
Result of sensitivity analysis of the 15 studies

### Determinant factors

#### Availability of assigned nurse in charge of individual care

Patients who had one nurse in charge of their care had 1.08 higher chance of being satisfied with nursing care compared to those patients without the assigned nurse in charge of their care although not statistically significant(OR: 1.08 (95% CI (0.45,2.62)) (Figure 6). The heterogeneity test (p=0.011) showed a significant evidence of variation across studies. Moreover, the result of Egger’s test showed no statistically significant evidence of publication bias (P=0.541).

**Figure 6:**
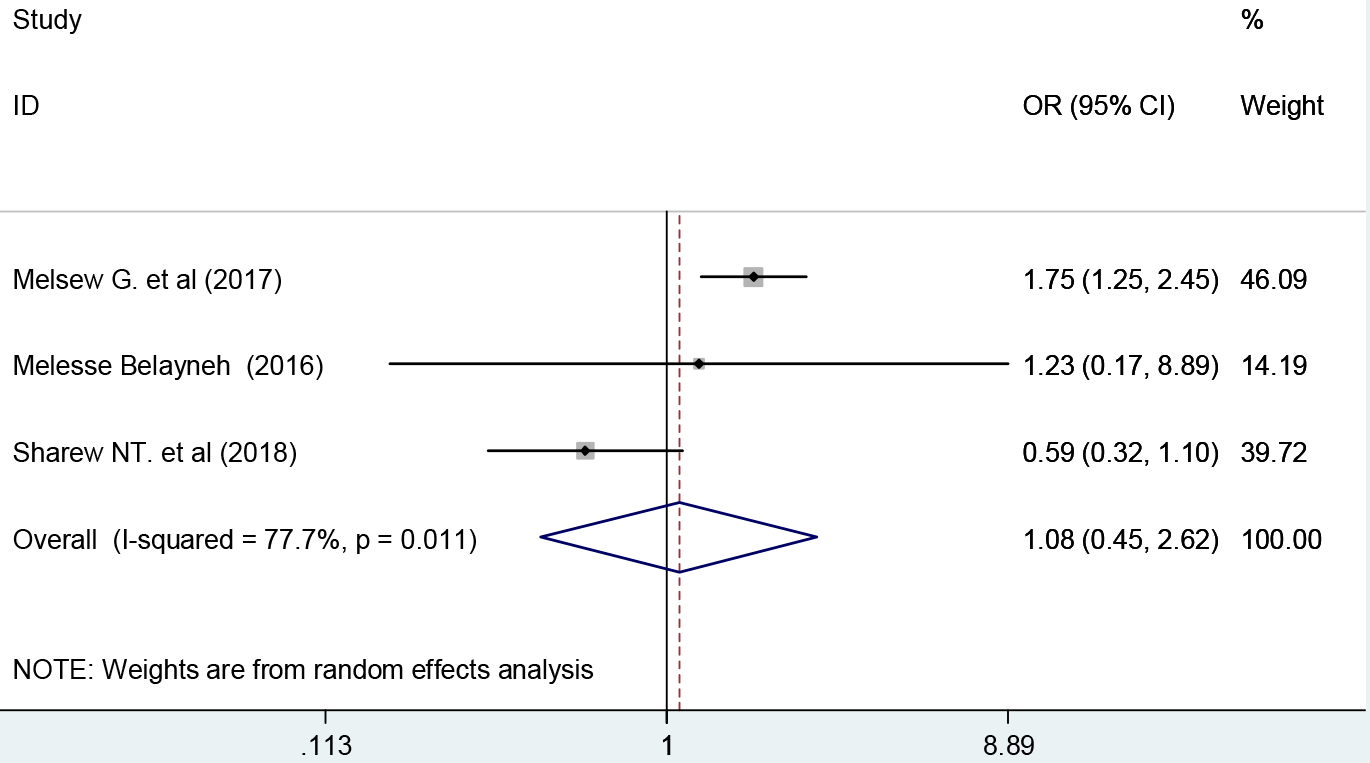
Forest plot showing the association between patient satisfaction and availability of one assigned nurse in charge of patient care

#### Place of residence

Patients living in the urban area had 1.07 higher chance of being satisfied with nursing care compared to those patient in a rural area although not statistically significant (OR: 1.07 (95% CI (0.70, 1.65)) (Figure 7). The heterogeneity test (P= 0.071) showed no significant evidence of variation across studies, Moreover, the result of Egger’s test showed a significant evidence of publication bias(P=0.012).

**Figure 7:**
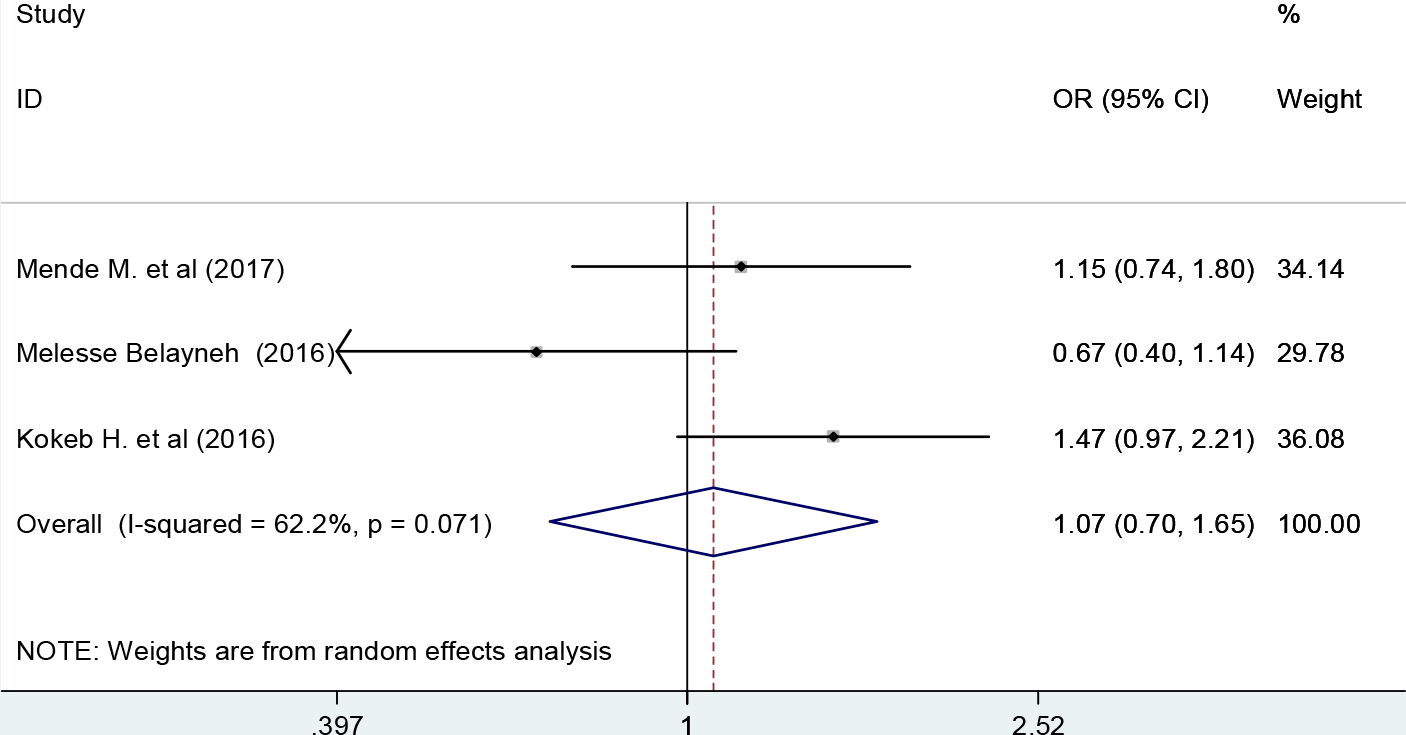
Forest plot showing the association between patient satisfaction and residence

#### History of admission

Patients who had no history of previous hospitalization had 1.37 higher chance of being satisfied with nursing care compared to those patients with a history of admission although not statistically significant (OR: 1.37 (95% CI (0.82,2.31)) (Figure 8). The heterogeneity test (P= 0. 001) showed a significant evidence of variation across studies. Moreover, the result of Egger’s test showed no statistically significant evidence of publication bias (P=0.25).

**Figure 8:**
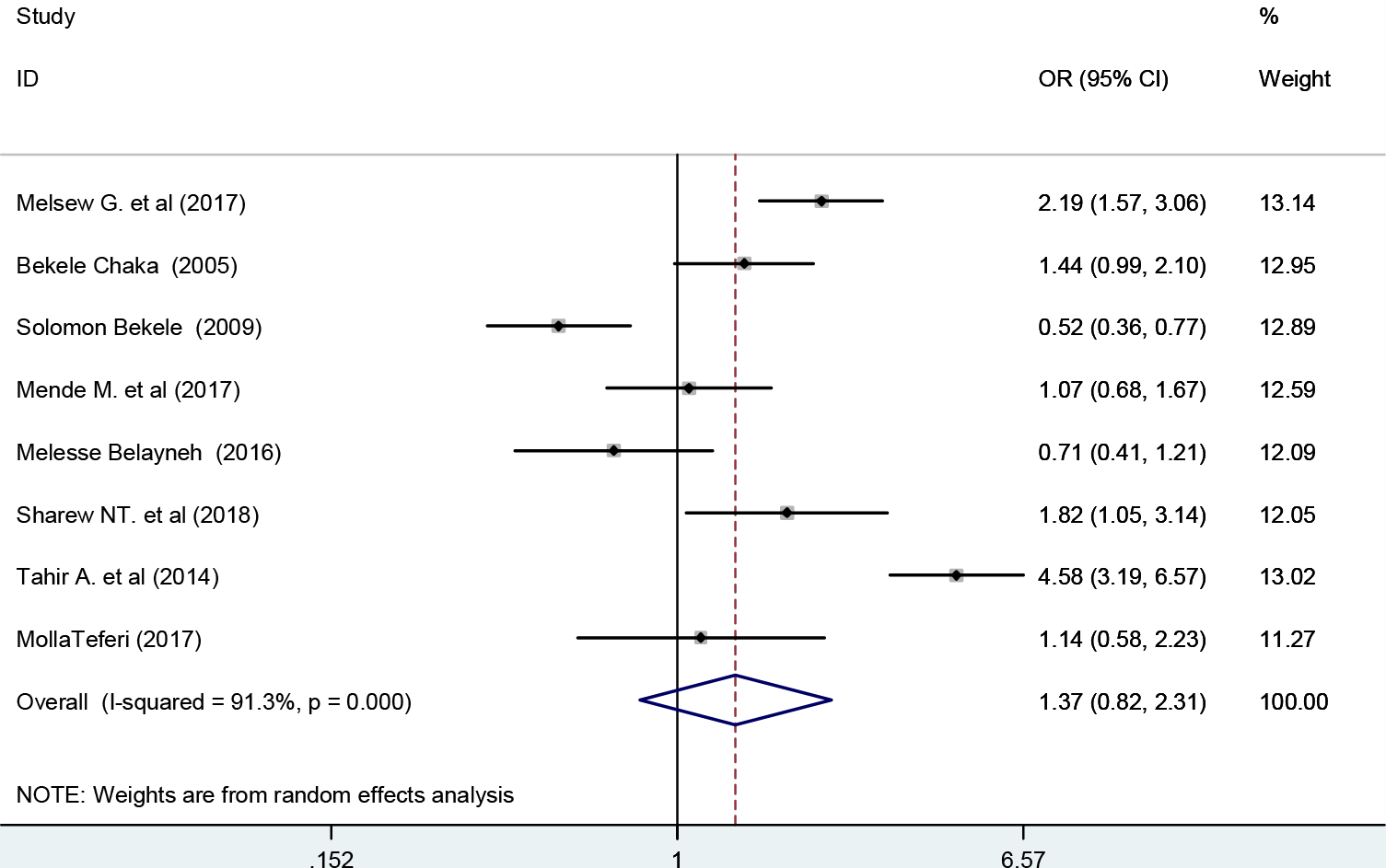
Forest plot showing the association between patient satisfaction and history of admission

#### Presence of other diseases

Patients who had no comorbid disease had 1.08 higher chance of being satisfied with nursing care compared to those patients without comorbidity (OR: 1.08 (95% CI (0.48, 2.39)) (Figure 9). The heterogeneity test showed a significant evidence of variation across studies, P= 0.001. Moreover, the result of Egger’s test to examine publication bias showed no statistically significant evidence of publication bias (P=0.91).

**Figure 9:**
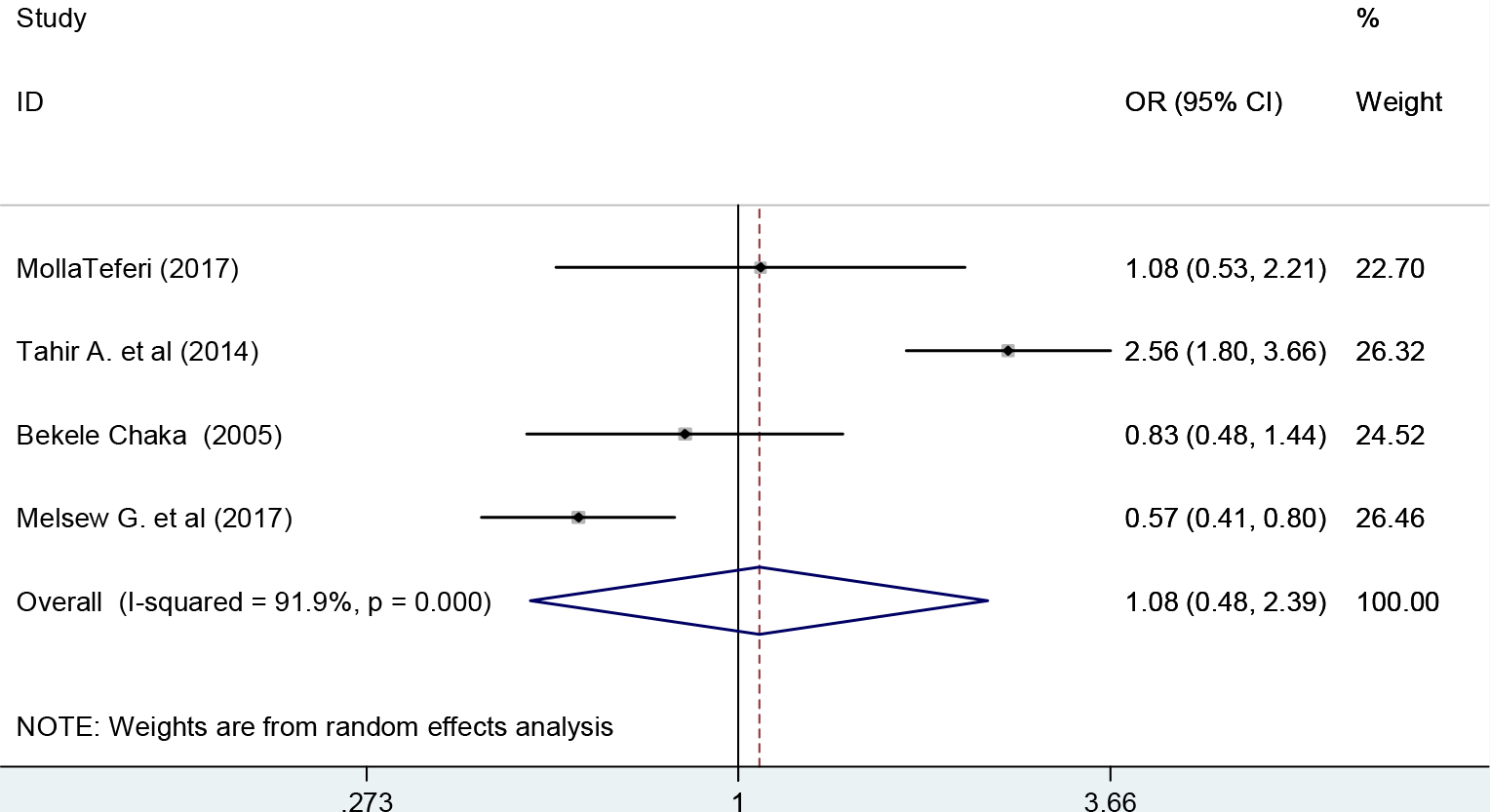
Forest plot showing the association between patient satisfaction and presence of other diseases

## Discussion

Within the healthcare of today, nurses spend more time by giving bedside nursing care for admitted patients than any other healthcare professionals in the hospital. Hence, patient satisfaction with nursing care is a definitive determinant of the quality of healthcare in the hospital [19, 48, 49]. In this systematic review and meta-analysis, we estimated the pooled proportion of satisfied patients with nursing care in Ethiopia.

Assessing the level of patient satisfaction with nursing care is crucial to improving the quality of care, also patient satisfaction has been considered as an indicator of patient outcome[50]. Moreover, patients’ overall satisfaction has a positive correlation with the health care provided at the hospital[51].

The result of this meta-analysis revealed that the overall estimated pooled level of patient satisfaction with nursing care was 55.15%. This finding was similar to previous studies conducted in Serbia and the Philippines in which 51.7% and 57.8% of patients were satisfied with nursing care respectively[18, 52].

The level of patient satisfaction with nursing care in our study was lower than other similar studies report of 69% in Iran[22], 67% in Kenya[23], 69.4% in Jordan[53], 82.7% in Malaysia[5], and 89.6% in Saudi Arabia[7]. This could be due to poor job satisfaction among Ethiopian nurses, low level of health care service and not well-qualified healthcare professionals in the country.

On the other hand, the level of patient satisfaction with nursing care in this study was higher than a study conducted in Ghana which revealed that about 33% of patients were satisfied with their nursing care[24]. Similarly, our finding was higher than other study conducted in public hospitals of Mosul City to assess patient satisfaction with nursing care which revealed 40% in ibn-Sina, 47% in Al-Jamhory, 42% in Al-Salam and 49% in AL-Kanssa teaching hospital[54]. The difference might be due to variation in sociodemographic characteristics of the study participants, sample size, measurement tool used to quantify the level of satisfaction, and others.

The result of this meta-analysis has found that patients’ residence, availability of assigned nurse in charge, previous history of admission, and the presence of other diseases had an influence on the patients’ satisfaction with nursing care even though not statistically significant. A similar studies revealed a significant association between patient satisfaction and previous admission to hospital[54, 55]. Poor quality of care, repeated costs, and bad experience during their past admission might be the possible reasons for patients with a history of admission to be dissatisfied with nursing care. Similar to our finding, a study done in England showed that availability of nurse in charge increases patients level of satisfaction with nursing care[56]. The possible reason might be due to the fact that patients could get a quick response from the available nurse for their needs and demand. Moreover, in our study urban patients were more satisfied than rural patients. This is in agreement with a recent systematic review[57].

Even though this review has provided valuable information and best evidence regarding the level of patient satisfaction with nursing care, there were some limitations, which we address below. First, our overall estimates showed significant heterogeneity among studies, so that interpretation of the results has to be taken cautiously. Although we performed subgroup analysis and metaregression, we could not identify the sources of variability. Second, it was difficult to analyze some additional major factors since they were not examined in a similar fashion across the studies. Third, it was difficult to compare our results with others due to lack of other published systematic review and meta-analysis on patient satisfaction with nursing. Finally, publication bias was appreciated even though it is inevitable in any meta-analysis.

### Conclusions and recommendations

The overall level of patient satisfaction with nursing was relatively moderate. Patient satisfaction was influenced by patients’ history of admission, residence, availability of assigned nurse, and presence of other diseases even though not statistically significant. This systematic review and meta-analysis provided a national evidence on the level of patient satisfaction with nursing care in Ethiopia. This might be very useful for policymakers to give more emphasis to the quality of nursing care in order to improve the overall quality of healthcare service. Furthermore, the Federal Ministry of health and Hospital administrator should give a great attention to the importance of quality nursing care to increase patient satisfaction.

## Abbreviations

CI: Confidence Interval
OR: Odds Ratio
PRISMA: Preferred Reporting Items for Systematic Reviews and Meta-Analyses
SNNP: Southern Nations, Nationalities, and Peoples
WHO: World Health Organizatino

## Declarations

### Consent for publication

Not applicable.

### Availability of data and materials

The data analyzed during the current systematic review and meta-analysis is available from the corresponding author on reasonable request.

### Ethics approval and consent to participate

Not applicable.

### Authors’ contributions

HM and GD developed the protocol and involved in the design, selection of study, data extraction, and statistical analysis and developing the initial drafts of the manuscript. FW, TDH and HB involved in data extraction, quality assessment, statistical analysis and revising subsequent drafts. HM and TDH prepared the final draft of the manuscript. All authors read and approved the final draft of the manuscript.

## Acknowledgment

We would like to thank all authors of studies included in this systematic review and metaanalysis.

## References

1. Kulkarni M, Dasgupta S, Deoke A, Nayse N: Study of satisfaction of patients admitted in a tertiary care hospital in Nagpur. National J Community Med 2011, 2(1):37–39.

2. Salmani N, Abbaszadeh A, Rasouli M, Hasanvand S: The process of satisfaction with nursing care in parents of hospitalized children: a grounded theory study. Int J Pediatr 2015, 3.

3. Tzeng H-M, Ketefian S, Redman RW: Relationship of nurses’ assessment of organizational culture, job satisfaction, and patient satisfaction with nursing care. International journal of nursing studies 2002, 39(1):79–84.

4. Ammo MA, Abu-Shaheen AK, Kobrosly S, Al-Tannir MA: Determinants of patient satisfaction at tertiary care centers in Lebanon. Open Journal of Nursing 2014, 4(13):939.

5. Teng K, Norazliah S: Surgical patients’ satisfaction of nursing care at the orthopedic wards in hospital universiti sains malaysia (HUSM). J Environ Health 2012, 3(1):36–43.

6. Aharony L, Strasser S: Patient satisfaction: what we know about and what we still need to explore. Medical care review 1993, 50(1):49–79.

7. Aldaqal SM, Alghamdi H, AlTurki H, El-deek BS, Kensarah A: Determinants of patient satisfaction in the surgical ward at a University Hospital in Saudi Arabia. Life Science Journal 2012, 9(1):277–280.

8. Wagner D, Bear M: Patient satisfaction with nursing care: a concept analysis within a nursing framework. Journal of advanced nursing 2009, 65(3):692–701.

9. Birhanu Darega, Nagasa Dida, Teshale Letimo, Tolashi Hunde, Yadashi Hayile, Shewafere Yeshitla, Amare M: Perceived Quality of nursing Cares Practices in Inpatient Departments of Bale Zone Hospitals, Oromiya Regional State, Southeast Ethiopia Facility-Based Cross Sectional Study. Quality in primary care 2016, 24(1):39–45.

10. Hughes F: Nurses at the forefront of innovation. International Nursing Review 2006, 53(2):94–101.

11. Batbaatar E, Dorjdagva J, Luvsannyam A, Savino MM, Amenta P: Determinants of patient satisfaction: a systematic review. Perspectives in public health 2017, 137(2):89–101.

12. Ahmed T, Assefa N, Demisie A, Kenay A: Levels of adult patients’ satisfaction with nursing care in selected public hospitals in Ethiopia. International journal of health sciences 2014, 8(4):371.

13. Tang WM, Soong C-Y, Lim WC: Patient satisfaction with nursing care: a descriptive study using interaction model of client health behavior. International Journal of Nursing Science 2013, 3(2):51–56.

14. Tsegaw MG: IN-PATIENTS’SATISFACTION LEVEL TOWARDS NURSING CARE SERVICES AND ASSOCIATED FACTORS AT PUBLIC HOSPITALS OF ADDIS ABABA, ETHIOPIA. Journal of Health, Medicine and Nursing 2017, 1(1): 1–17.

15. Ebrahim SM, Issa SS: Satisfaction with nursing care among patients attending oncology center in Basra city, Iraq. Journal of Environmental Science and Engineering 2015:241–248.

16. Joolaee S, Hajibabaee F, Jalal EJ, Bahrani N: Assessment of Patient Satisfaction from Nursing Care in Hospitals of Iran University of Medical Sciences. Hayat 2011, 17(1).

17. Larrabee JH, Ostrow CL, Withrow ML, Janney MA, Hobbs Jr GR, Burant C: Predictors of patient satisfaction with inpatient hospital nursing care. Research in Nursing & Health 2004, 27(4):254–268.

18. Milutinović D, Simin D, Brkić N, Brkić S: The patient satisfaction with nursing care quality: the psychometric study of the Serbian version of PSNCQ questionnaire. Scandinavian journal of caring sciences 2012, 26(3):598–606.

19. Mohammed HA, Ali RI, Mussa YM: Assessment of Adult Patients Satisfaction Regarding Nursing Care in Different Hospitals in Kirkuk City. kirkuk university journal for scientific studies 2016, 11(3):222–236.

20. Freitas JSd, Silva AEBdC, Minamisava R, Bezerra Alq, Sousa MRGd: Quality of nursing care and satisfaction of patients attended at a teaching hospital. Revista latino-americana de enfermagem 2014, 22(3):454–460.

21. Zahr LK, William SG, Ayam E-H: Patient satisfaction with nursing care in Alexandria, Egypt. International journal of nursing studies 1991, 28(4):337–342.

22. Farahani MF, Shamsikhani S, Hezaveh MS: Patient satisfaction with nursing and medical care in hospitals affiliated to arak university of medical sciences in 2009. Nursing and midwifery studies 2014, 3(3).

23. Ndambuki J: The level of patients’ satisfaction and perception on quality of nursing services in the Renal unit, Kenyatta National Hospital Nairobi, Kenya. Open Journal of Nursing 2013, 3(02):186.

24. Dzomeku V, Ba-Etilayoo A, Perekuu T, Mantey R: In-patient satisfaction with nursing care: a case study at Kwame Nkrumah University of Science and Technology hospital. International Journal of Research in Medical and Health Sciences 2013, 2(1):19–24.

25. Samina M, Qadri G, Tabish S, Samiya M, Riyaz R: Patient’s perception of nursing care at a large teaching hospital in India. International Journal of Health Sciences 2008, 2(2):92.

26. Belayneh M: Inpatient satisfaction and associated factors towards nursing care at Felegehiwot Referral Hospital, Amhara Regional State, Northwest Ethiopia. GLOBAL JOURNAL OF MEDICINE AND PUBLIC HEALTH 2016, 5(3).

27. Mensa M, Taye A, Katene S, Abera F, Ochare O: Determinants of Patient Satisfaction Towards Inpatient Nursing Services and its Associated Factors in, Gamo Gofa Zone, SNNPR, Ethiopia, April 2017. MOJ Clin Med Case Rep 2017, 7(3):00205.

28. Molla M, Berhe A, Shumye A, Adama Y: Assessment of adult patients satisfaction and associated factors with nursing care in Black Lion Hospital, Ethiopia; institutional based cross sectional study, 2012. International Journal of Nursing and Midwifery 2014, 6(4):49–57.

29. Liberati A, Altman DG, Tetzlaff J, Mulrow C, Gøtzsche PC, Ioannidis JP, Clarke M, Devereaux PJ, Kleijnen J, Moher D: The PRISMA statement for reporting systematic reviews and meta-analyses of studies that evaluate health care interventions: explanation and elaboration. PLoS medicine 2009, 6(7):e1000100.

30. Munn Z, Moola S, Riitano D, Lisy K: The development of a critical appraisal tool for use in systematic reviews addressing questions of prevalence. International journal of health policy and management 2014, 3(3):123.

31. Modesti PA, Reboldi G, Cappuccio FP, Agyemang C, Remuzzi G, Rapi S, Perruolo E, Parati G: Panethnic differences in blood pressure in Europe: a systematic review and meta-analysis. PLoS One 2016, 11(1):e0147601.

32. Higgins JP, Thompson SG: Quantifying heterogeneity in a metaanalysis. Statistics in medicine 2002, 21(11):1539–1558.

33. Ried K: Interpreting and understanding meta-analysis graphs: a practical guide. 2006.

34. McFarland LV: Meta-analysis of probiotics for the prevention of antibiotic associated diarrhea and the treatment of Clostridium difficile disease. The American journal of gastroenterology 2006, 101(4):812.

35. Teshome HM, Ayalew GD, Shiferaw FW, Leshargie CT, Boneya DJ: The Prevalence of Depression among Diabetic Patients in Ethiopia: A Systematic Review and Meta-Analysis, 2018. Depression Research and Treatment 2018, 2018.

36. Egger M, Smith GD, Phillips AN: Meta-analysis: principles and procedures. Bmj 1997, 315(7121):1533–1537.

37. Tura G, Fantahun M, Worku A: The effect of health facility delivery on neonatal mortality: systematic review and meta-analysis. BMC pregnancy and childbirth 2013, 13(1):18.

38. al GGe: WHY PATIENTS ARE DISSATISFIED ON NURSING CARE SERVICES AT MENELIK HOSPITAL, ADDIS ABABA. Journal of Bio Innovation 2016, 5(6).

39. Bekele S: Patient Satisfaction with Nursing Care in Medical and Surgical Wards of government hospitals, Addis Ababa, Ethiopia. 2009.

40. Chaka B: Adult patient satisfaction with nursing care. MPH thesis, department of community health, Addis Ababa University 2005.

41. Alemu S, Jira C, Asseffa T, Desa MM: Changes in inpatient satisfaction with nursing care and communication at Debre Markos Hospital, Amhara Region, Ethiopia. American Journal of Health Research 2014, 2(4):171–176.

42. Haile Eyasu K, Adane AA, Amdie FZ, Getahun TB, Biwota MA: Adult Patients’ Satisfaction with Inpatient Nursing Care and Associated Factors in an Ethiopian Referral Hospital, Northeast, Ethiopia. Advances in Nursing 2016, 2016.

43. Negash AK, Negussie D, Demissie AF: Patients’ satisfaction and associated factors with nursing care services in selected hospitals, Northwest Ethiopia. American Journal of Nursing Science 2014, 3(3):34–42.

44. Sharew NT, Bizuneh HT, Assefa HK, Habtewold TD: Investigating admitted patients’ satisfaction with nursing care at Debre Berhan Referral Hospital in Ethiopia: a cross-sectional study. BMJ open 2018, 8(5):e021107.

45. Legesse M, Salgedo W, Walle A: Adult patient satisfaction with in-patient nursing care in a Referral and Teaching Hospital in Southern Nations Nationalities and Peoples’ Region (SNNPR), Ethiopia. Journal of Nursing and Care 2016, 5(334):2167–1168.

46. Jiru TG: Adult in Patients’ Satisfaction and Associated Factors with Nursing Care in Wards of Hospitals of in Guji Zone, Oromia, South Ethiopia. Journal of Medical Reviews 2018, 1(1):1–10.

47. MollaTeferi: Assessment of Adult Patients’ Perception of Nursing Care and Its Contributing Factors at Ayder Referral Hospital, Mekelle, Ethiopia. Journal of Pharmacy and Alternative Medicine 2017, 14.

48. González-Valentín A, Padín-López S, de Ramón-Garrido E: Patient satisfaction with nursing care in a regional university hospital in southern Spain. Journal of Nursing Care Quality 2005, 20(1):63–72.

49. Uzun Ö: Patient satisfaction with nursing care at a university hospital in Turkey. Journal of nursing care quality 2001, 16(1):24–33.

50. Cleary PD, McNeil BJ: Patient satisfaction as an indicator of quality care. Inquiry 1988:25–36.

51. Al-Mailam FF: The effect of nursing care on overall patient satisfaction and its predictive value on return-to-provider behavior: a survey study. Quality Management in Healthcare 2005, 14(2):116–120.

52. Villarruz-Sulit MVC, Dans AL, Javelosa MAU: Measuring satisfaction with nursing care of patients admitted in the medical wards of the philippine general hospital. Acta Medica Philippina 2009, 43(4):52–56.

53. Alasad JA, Ahmad MM: Patients’ satisfaction with nursing care in Jordan. International Journal of Health care quality assurance 2003, 16(6):279–285.

54. Mukhlif HH: Assessment of Adult Patients Satisfaction with the Nursing Care in the Public Teaching Hospitals of Mosul City. kufa Journal for Nursing sciences 2014, 4(2):34–41.

55. Findik UY, Unsar S, Sut N: Patient satisfaction with nursing care and its relationship with patient characteristics. Nursing & health sciences 2010, 12(2):162–169.

56. Aiken LH, Sloane DM, Ball J, Bruyneel L, Rafferty AM, Griffiths P: Patient satisfaction with hospital care and nurses in England: an observational study. BMJ open 2018, 8(1):e019189.

57. Salehi A, Jannati A, Nosratnjad S, Heydari L: Factors influencing the inpatients satisfaction in public hospitals: a systematic review.

